# Inhibition of phosphodiesterase 4B as a novel therapeutic strategy for the treatment of refractory epilepsy

**DOI:** 10.1101/2025.05.13.653835

**Authors:** Cassandra D. Kinch, Mitchell Kesler, Deepika Dogra, Kyle Morin, Kingsley Ibhazehiebo, Christie G. Noschang, Jong M. Rho, Deborah M. Kurrasch

## Abstract

Despite the availability of nearly 40 approved anti-seizure medications (ASMs), at least one-third of individuals with epilepsy remain refractory to treatment, and many experience life-limiting cognitive or psychiatric side effects. Using a machine learning-guided platform, we identified PDE4 as an underexplored anti-seizure target, which became further validated based on its enriched expression in seizure-relevant brain regions and its potential to modulate excitatory/inhibitory neuronal tone via cAMP signaling. The pan-PDE4 inhibitor crisaborole partially protected against hyperthermia-induced seizures and reduced spontaneous seizures in *Scn1a+/-* mice, while rolipram and roflumilast showed no efficacy at tolerable doses. SN-2000, a first-in-kind allosteric modulator of PDE4B, was rationally designed for isoform selectivity and brain penetration, and demonstrated versatile reduction of seizure activity across multiple zebrafish and rodent genetic and acquired epilepsy models, with efficacy comparable to standard-of-care ASMs. SN-2000 also demonstrated favorable behavioral outcomes, reducing post-ictal aggression and anxiety-like behaviors, and improving cognitive performance in both wild-type and epileptic mice. These effects were linked to paradoxical regulation of excitatory and neuronal activity in the cortex and thalamus of epileptic mice, respectively, as well as elevated cAMP signaling and downstream pCREB activation. Together, these findings support PDE4B inhibition as a disease-relevant mechanism in epilepsy, and position SN-2000 as a promising therapeutic candidate offering seizure control without the neuropsychiatric burden of existing ASMs and potential pro-cognitive properties.

**One Sentence Summary:** SN-2000, a novel allosteric PDE4 inhibitor, reduces seizure activity and shows psychiatric-neutral and pro-cognitive properties in preclinical models

## INTRODUCTION

Epilepsy is the fourth most prevalent neurological disorder globally, with an estimated 50 million individuals affected (Feigin et al. 2019). Its incidence is surpassed only by stroke, neurodegenerative dementias (including Alzheimer’s disease), and Parkinson’s disease. Despite 100 years of epilepsy drug discovery and nearly 40 approved ASMs, most patients with epilepsy fail to respond to first- and second-line therapies. Even across epileptic patients whose seizures are controlled by ASMs, nearly half report life-limiting side effects, including cognitive and psychiatric dysfunction (Fattorusso et al. 2021, Gilliam et al. 2002). Thus, drugs with novel mechanisms of action that reduce seizure activity while limiting treatment-emergent adverse events (TEAEs) are desperately needed.

Using our machine learning-derived platform technology designed to reveal unappreciated druggable epilepsy targets, we uncovered phosphodiesterase 4B inhibition (PDE4Bi) as promising strategy (Ibhazehiebo et al 2018). PDE4 is an intracellular enzyme that catabolizes cyclic adenosine 3’,5’-monophosphate (cAMP) (derived from adenosine triphosphate) to 5’adenosine monophosphate (Blockland et al. 2019). The PDE4 family has four isoforms – PDE4A, 4B, 4C, and 4D – which differ in structure due to their N-terminal regulatory domains, as well as tissue expression and subcellular localization (Donders et al 2024, Baillie et al. 2020).

Although underappreciated in epilepsy, two studies link increased cAMP with seizure termination, likely due to neuronal stabilization in ictal brain areas (Ferrendelli et al. 1980, Yang et al. 2024). Mechanistically, increases in cAMP elevates protein kinase A (PKA) and exchange protein directly activated by cAMP (EPAC), which may increase inhibitory tone through enhanced GABA vesicle recycling (Donders et al, Nakamura et al 2015). Concomitantly, cAMP interacts with hyperpolarization-activated cyclic nucleotide-gated channels (HCNs) and cyclic nucleotide-gated channels (CNGCs) to depolarize glutamatergic cells, potentially increasing excitatory firing thresholds (Donders et al, Huang and Kandel 1996, Lyman et al 2021). Beyond the immediate impact on excitatory/inhibitory signaling, PDE4i, through increased cAMP, can modulate expression of GABA-A1R, GluA1-AMPA, BDNF, CaMK/p38 MAPK, and NF-kB, resulting in enduring effects on inhibitory tone, excitotoxicity, synaptic plasticity, neuronal survival, and neuroinflammation, respectively (Nakamura et al 2015, Donders et al 2025).

Phase 1/2 trials with PDE4i demonstrated clinically relevant cognitive improvements in healthy elderly adults, as well as Schizophrenic and Alzheimer’s disease patients (Sugin et al. 2020, Crocetti et al. 2022). Moreover, Phase 2/3 trials demonstrated efficacy of PDE4i in major depressive disorder (Donders et al 2025, Crpcetto et al 2022). However, clinical development of these older generation pan-PDE4is was untenable due to TEAEs involving emesis, nausea and diarrhea (Donders et al. 2025). To circumvent tolerability issues of pan-PDE4is, the newer generations of PDE4is are rationally designed to achieve isoform specificity through exploitation of isoform-specific structural differences in the active site, or allosteric binding at regulatory sites on the PDE4 protein. Indeed, a Phase 2 trial in adults with Fragile X syndrome (FXS) reported comparable tolerability and safety with the PDE4D allosteric inhibitor zatomilast versus placebo (Berry-Kravis et al. 2021). Recruitment is ongoing for registrational trials of zatomilast in adolescents and adults with FXS (NCT05358886, NCT05163808, NCT05367960).

Here, we report proof-of-concept data supporting the role for PDE4 inhibition in reducing seizure frequency in a severe, refractory epilepsy model. We also report development of a first-in-class isozyme specific allosteric modulator of PDE4Bi, SN-2000, and provide seminal evidence of the compounds’ preclinical efficacy in reduction of seizure frequency, as well as psychiatric-neutral and potential pro-cognitive properties.

## RESULTS

### PDE4is stabilize neurons and provide partial seizure protection

As an initial proof-of-concept, we tested the protective capabilities of three commercially available PDE4is against febrile seizures in the well established, refractory epileptic *Scn1a+/-* mouse model (Hawkins et al. 2017). While acute rolipram and roflumilast resulted in only 10% (1/10) and 0% (0/4) protection from hyperthermia-induced seizures, acute crisaborole treatment resulted in 64% (6/11) protection, and was statistically significant vs placebo (p=0.0475, Figure 1a). Additionally, the number of spontaneous recurrent seizures in *Scn1a+/-* mice was reduced in crisaborole vs vehicle treated mice at all three treatment days, with seizure freedom achieved on day 2 and 3 (Figure 1b). To confirm the restorative ability of crisaborole on cAMP signaling in epileptic models, we sectioned cortexes of spontaneously seizing *Scn1a+/-* mice treated with crisaborole vs vehicle and observed cytoplasmic relocation of pCREB suggestive of PKA activation (Figure 1c). Cortical sections from crisaborole treated mice also showed a dampening of excitatory neuronal activity, while thalamic sections from the same mice showed increased inhibitory neuronal activity, supporting the hypothesized paradoxical inhibitory/excitatory role of PDE4i in stabilizing neuronal tone in epileptic brains (Figure 1d,e).

**Figure 1:**
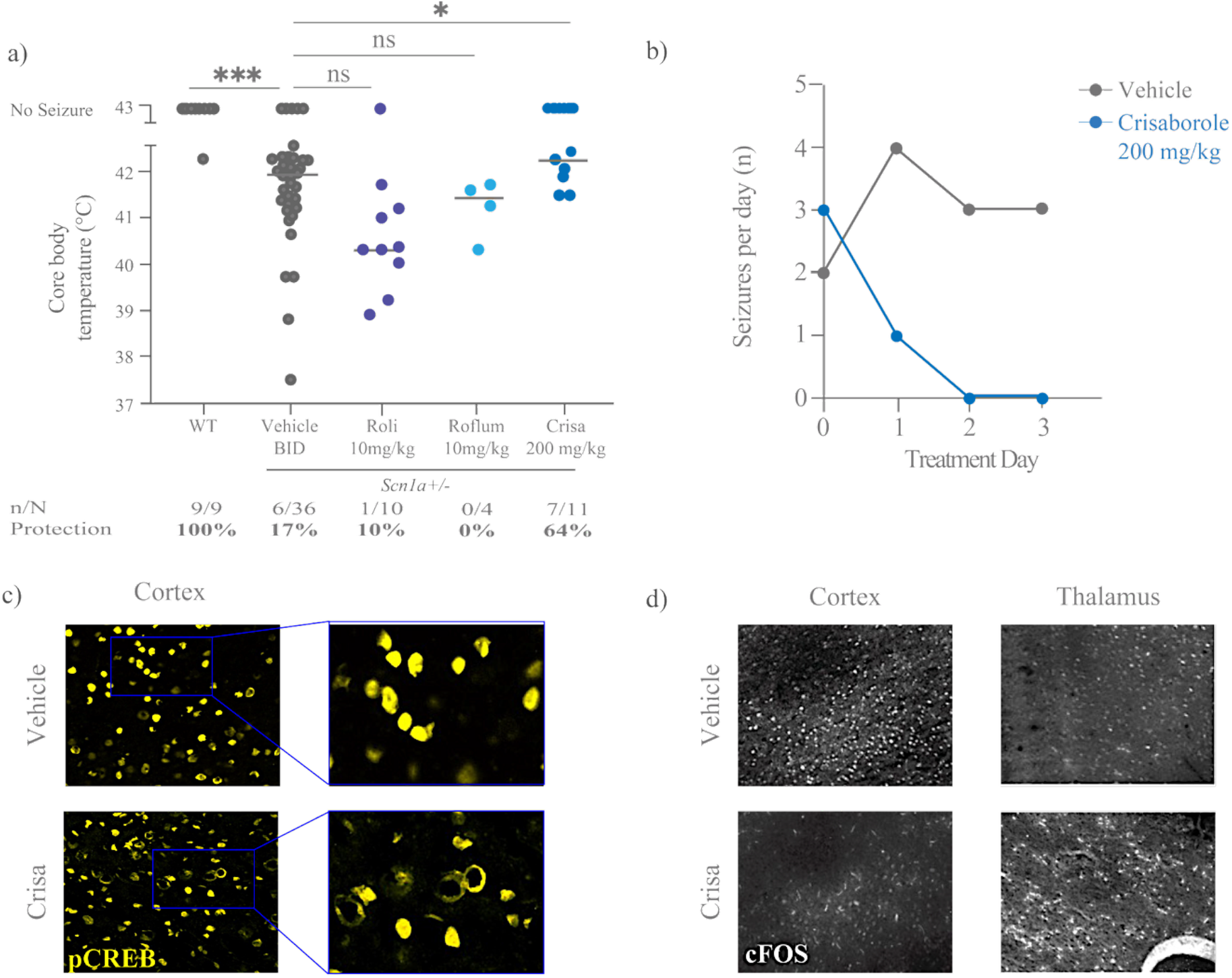
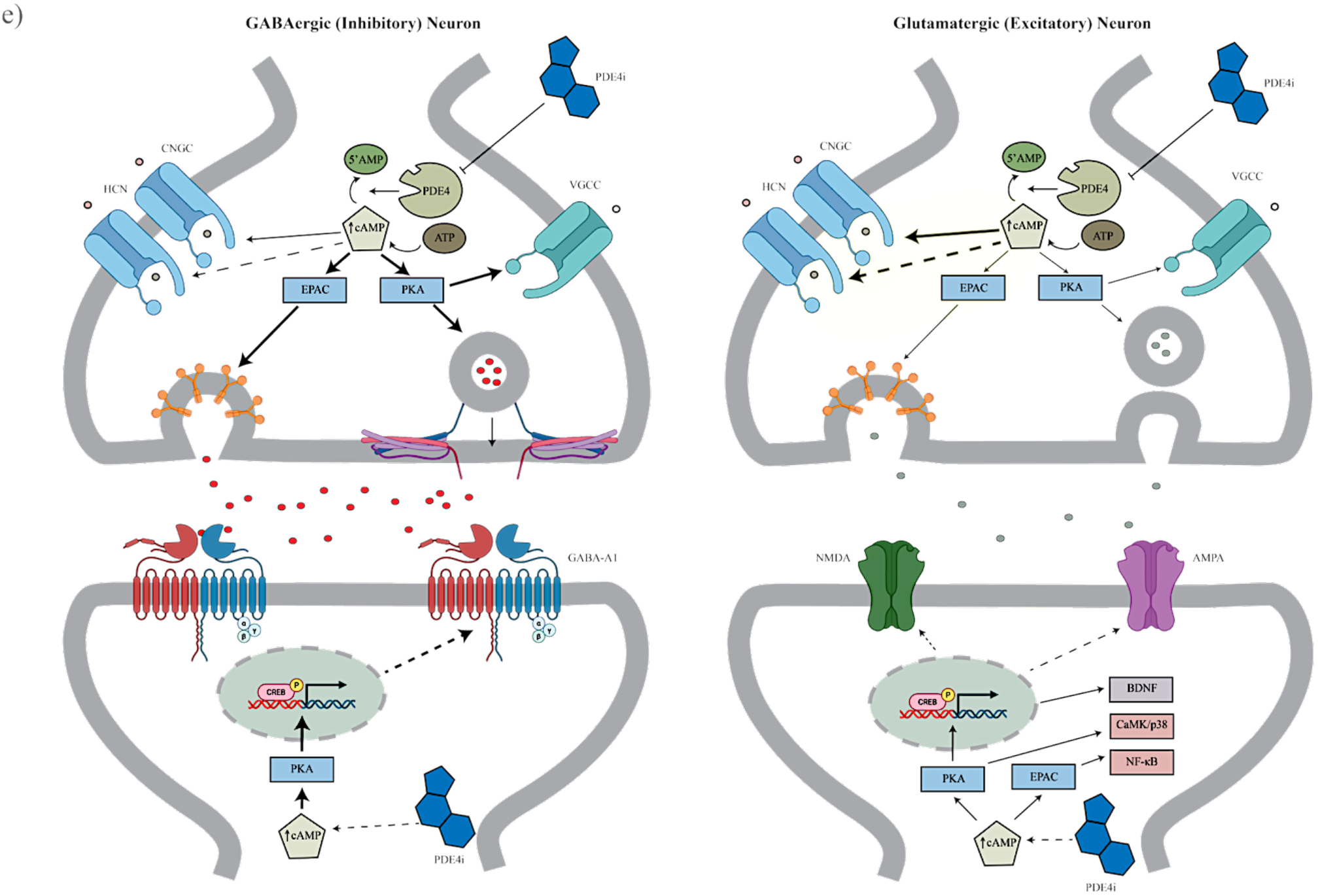
PDE4i exposure protects against seizures in refractory epilepsy model (*Scn1a+/-*) through restoration of inhibitory and excitatory neuronal stabilization. a) Protection of hyperthermia-induced seizures in *Scn1a+/-* mice treated with the commercially available PDE4is rolipram 10mg/kg, roflumilast 10mg/kg, and crisaborole 200mg/kg, or vehicle. b) Number of seizures per day recorded via video EEG (vEEG) in spontaneously seizing *Scn1a+/-* epileptic mice exposed to crisaborole 200mg/kg or vehicle. c) pCREB expression in cortical slices of *Scn1a+/-* mice exposed to crisaborole 200mg/kg or vehicle. d) Neuronal activation (cFOS) in cortical and thalamic slices of *Scn1a+/-* mice exposed to vehicle or crisaborole 200mg/kg. e) Schematic diagram of proposed mechanism of PDE4i-mediated reduction in seizure activity through regulation of inhibitory and excitatory neuronal transmission. Kruskal-Wallis test (p<0.0001) with Dunn’s multiple comparisons test, ***p<0.0001, *p<0.05. BDNF=brain-derived neutrophic factor; CaMK/p38= calcium/calmodulin-dependent protein kinase; cFOS=cellular homolog of the Finkel-Biskis-Jinkins osteosarcoma viral oncogene; Crisa=crisaborole; CNGC=cyclic nucleotide-gated channels; EPAC=exchange protein directly associated with cAMP; HCN=hyperpolarization-activated cyclic nucleotide-gated channel; ns=not significant; NF-kB=neutrophic factor-kB; pCREB=phosphorylated cAMP response element binding protein; PDE4i=phosphodiesterase 4 inhibitor; PKA=protein kinase A; Roflum=roflumilast; Roli=rolipram; VGCC=voltage-gated calcium channel

### SN-2000, a first-in-class allosteric PDE4B inhibitor for reduction of seizure activity

Considering the tolerability issues characteristic of older generation pan-PDE4is, we initiated development of a PDE4 isoform-specific inhibitor strategy. Of the four PDE4 isoforms (PDE4A-D), PDE4B shows broadest expression in brain regions associated with seizure-genesis, and less associated with emetic centers (Lamontagne et al 2001, Richter et al. 2013, Perez-Torrez et al., Burgin et al). To characterize association of specific PDE4 isoforms in modulating seizure activity, agarose-embedded *scn1lab-/-* epileptic zebrafish embryos (*Danio rerio*) were injected with knock-down morpholinos targeting *pde4a-d* (a and b zebrafish-specific orthologs were included) and monitored for seizure-like activity via extracellular field recordings as previously described (Ibhazeheibo et al. 2018). Seizure-like activity was reduced approximately 4-fold in PDE4B-MO and PDE4C-MO injected zebrafish, 3-fold in PDE4D-MO injected zebrafish and 1-fold in PDE4A-MO injected zebrafish compared to scramble-MO injected zebrafish (Figure 2a). With PDE4B confirmed as the target isoform, we designed SN-2000 as a novel brain-penetrant compound that binds allosterically at the dimerized PDE4 long form splice variant, yielding PDE4B isoform specificity (Figure 2b). Target engagement of SN-2000 at the PDE4B allosteric site vs the catalytic site, and at PDE4B vs PDE4D was confirmed through an in vitro competitive binding assays (Figure 2c,d). Last, intracellular activity of SN-2000, and A33, an established PDE4B inhibitor, was confirmed through a 3-fold increase of cAMP accumulation in cells overexpressing PDE4B vs PDE4A/D isoforms (Figure 2e).

**Figure 2.**
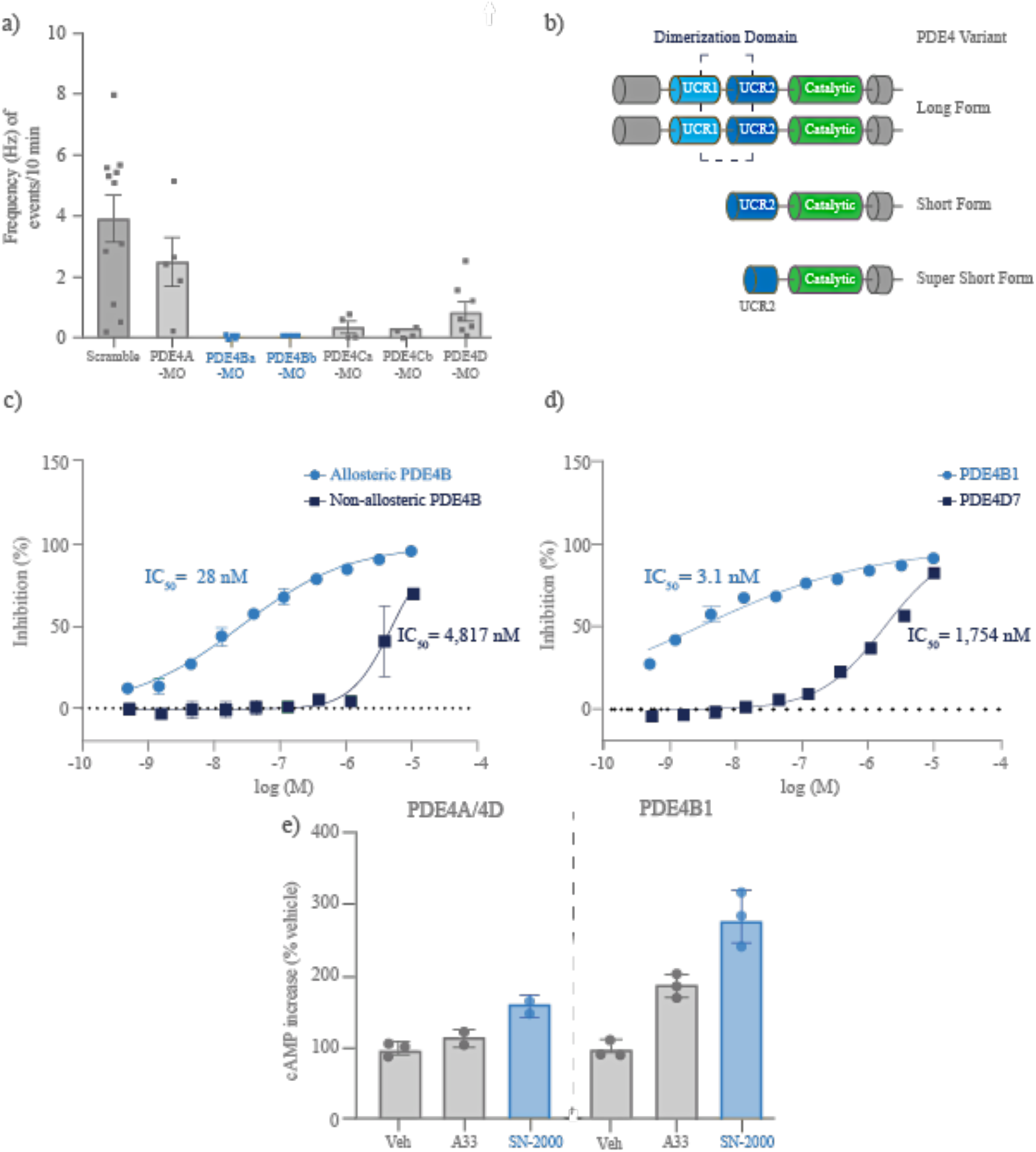
SN-2000 binds to a PDE4B allosteric site with nanomolar potency and selectivity over PDE4A/D. a) Frequency of seizure-like events in *scn1lab-/-* zebrafish injected with individual PDE4(A-D) isoform or scramble knockdown morpholinos. b) Cartoon diagram of PDE4B splice variants, including dimerized long form, short form and super short forms. C) Competitive inhibitory activity of SN-2000 at the allosteric PDE4B site vs non-allosteric (catalytic) site. c) Competitive inhibitory activity of SN-2000 at PDE4B vs PDE4D isoforms. e) Intracellular cAMP increase in cell lines overexpressing PDE4A/4D or PDE4B exposed to SN-2000, A33 (PDE4B inhibitor) or vehicle. MO=morpholino; UCR=upstream conserved region; Veh=Vehicle

### SN-2000 exposure confers protection from febrile and spontaneous recurrent seizures in refractory epilepsy model

We first evaluated anti-seizure properties of SN-2000 in the *scn1lab-/-* epileptic zebrafish model using our established locomotor seizure-like activity assay (Ibhazeheibo et al 2018).

Exposure to SN-2000 resulted in a significant decrease in seizure-like activity vs vehicle, which was comparable to first- and second-line ASMs (valproic acid [VPA] and VPA + clobazam [CLO]; Veh vs SN-2000, p<0.01; SN-2000 vs VPA, ns; SN-2000 vs VPA+CLO, ns; Figure 3a). In *Scn1a+/-* mice, SN-2000 demonstrated partial protection (75%) from hyperthermia-induced seizures, which was comparable to standard of care ASMs (VPA [69%] and CLO [88%]; Veh vs SN-2000 p<0.01; SN-2000 vs VPA, CLO, ns; Figure 3b). Further, a statistically significant reduction in spontaneous recurrent seizure activity was observed when comparing SN-2000 40mg/kg BID vs vehicle, measured by video EEG (p<0.0001, Figure 3c). Notably, vehicle exposed *Scn1a+/-* mice showed an abundance of interictal spike activity, which was not observed in SN-2000 40mg/kg exposed mice, indicative of a reduction of network hyperexcitability.

**Figure 3.**
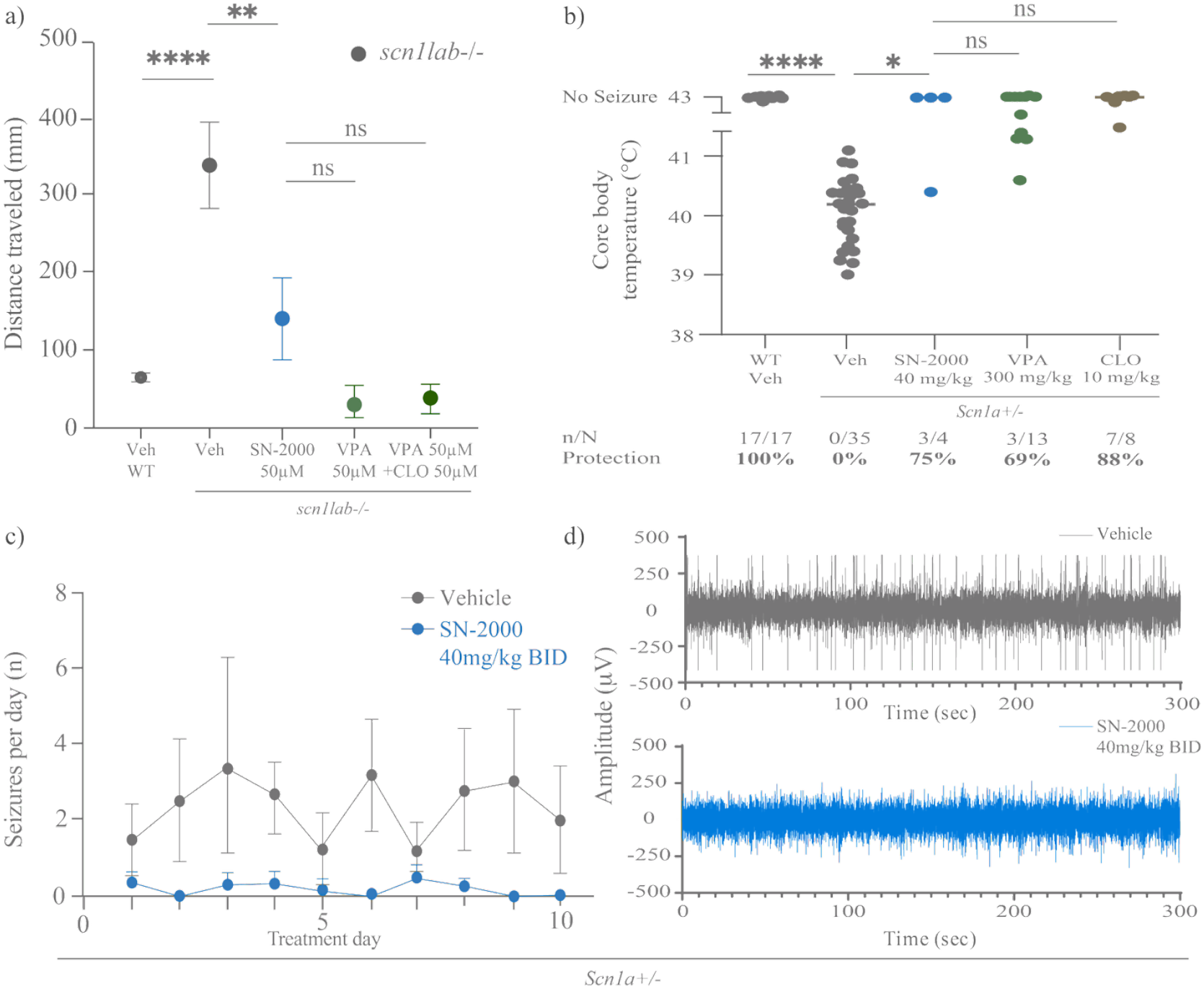
Reduction of seizure activity with SN-2000 is comparable to standard of care in refractory 152 epilepsy model. a) Reduction of seizure-like activity in *scn1lab-/-* epileptic zebrafish exposed to SN-2000, VPA, VPA+CLO or vehicle. b) Rate of protection from febrile seizures in *Scn1a+/-* epileptic mice exposed to SN-2000 40mg/kg, VPA 300mg/kg, CLO 10mg/kg, or vehicle. c) Number of seizures per day recorded via video EEG in *Scn1a+/-* epileptic mice exposed to SN-2000 40mg/kg or vehicle. Mice experiencing Racine level 4 seizures within the first 48 hours of treatment were excluded. d) Representative interictal tracing in *Scn1a+/-* mice exposed to SN-2000 40mg/kg or vehicle that were seizure free for ∼1 day. (a) One-way ANOVA p<0.0001, Sidak’s multiple comparisons test, ****p<0.0001, **p<0.01. (b) Kruskal-Wallis test (p<0.0001) with Dunn’s multiple comparisons test, ***p<0.0001, *p<0.05. (c) Mixed effects model (REML) Two-factor ANOVA, p=<0.0001. CLO=clobazam; ns=not significant; Veh=vehicle; VPA=valproic acid

### SN-2000 reduces seizure activity across genetic and acquired models of generalized epilepsy

To test the phenotype specificity of SN-2000 in reducing seizure activity we generated Kv1.1 (*kcn1a-/-*) channel and GABA_A1 (*gabra1-/-*) receptor knockout, and PTZ-induced epileptic zebrafish models (Dogra et al. 2023). Exposure to SN-2000 vs vehicle resulted in a significant reduction in seizure-like activity across all epileptic zebrafish models, which as comparable to standard of care ASMs (VPA, carbamazepine [CBZ], levetiracetam [LEV], lamotrigine [LTG] and VPA + CLO) for generalized epilepsy (Veh vs SN-2000, p<0.0001; SN-2000 vs VPA, ns; SN-2000 vs CBZ, ns; SN-2000 vs LEV, p<0.01; SN-2000 vs LTG, ns; SN-2000 vs VPA+CLO, ns; Figure 4a). Protection from maximal electroshock stimulated seizures was observed in 50% (n=2/4) of SN-2000 40mg/kg exposed wild type (WT) mice, 75% (n=6/8) of SN-2000 60mg/kg exposed WT mice, and 75% (n=6/8) of WT mice exposed to SN-2000 80mg/kg, which was comparable to protection observed in VPA exposed WT mice (88%). Together, these data suggest SN-2000 provides robust reduction of seizure activity across these genetic and acquired epilepsy models, with effect sizes comparable to standard of care ASMs.

**Figure 4:**
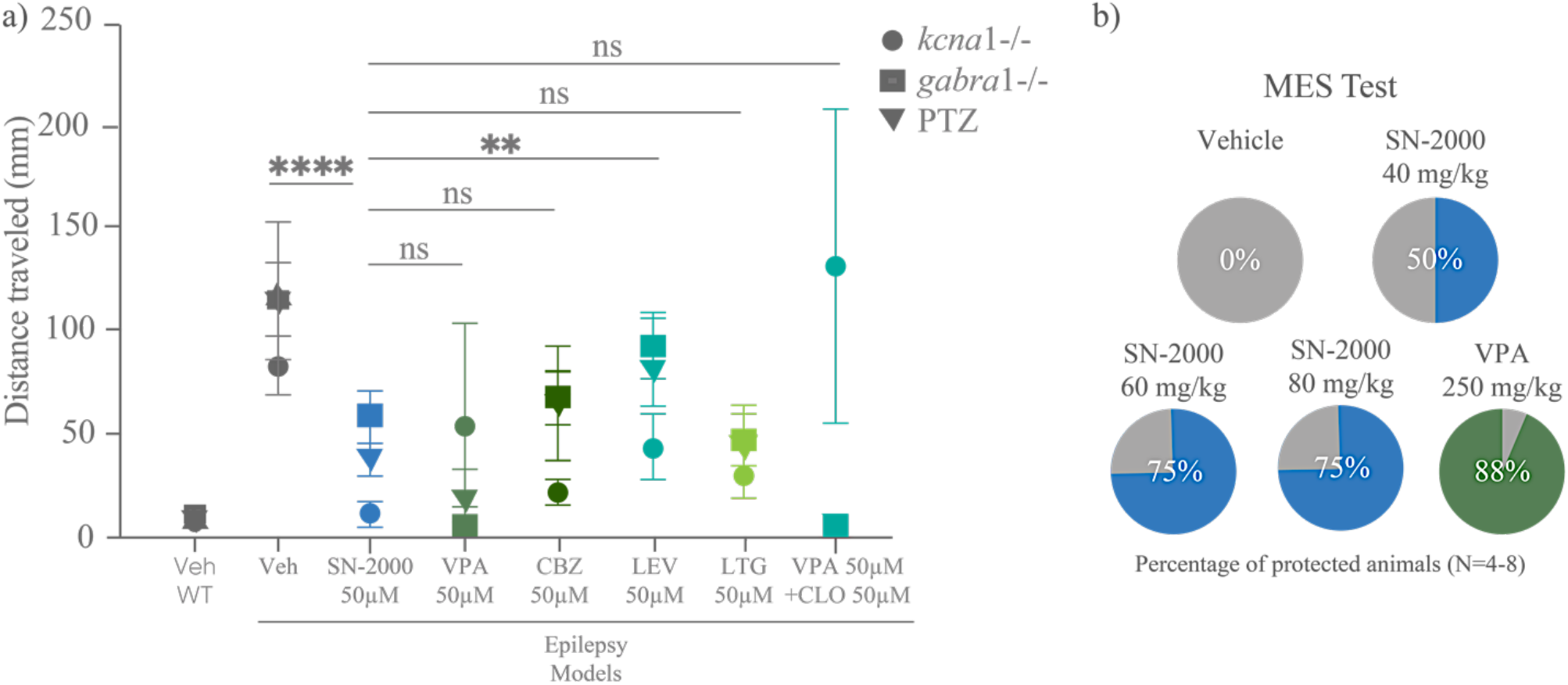
SN-2000 exposure reduces seizure activity across to levels comparable to SoC, across epilepsy models. a) Seizure-like activity in *kcna1-/-, gabra1-/-* or PTZ-induction zebrafish epilepsy models exposed to SN-2000, VPA, CBZ, LEV, LTG, VPA+CLO or vehicle. b) Protection from maximal electroshock (MES) seizure induction in SN-2000 40mg/kg, SN-2000 60mg/kg, SN-2000 80mg/kg or VPA exposed wild type (WT) mice. Two-way ANOVA, p<0.0001, Sidak’s multiple comparisons test, ****p<0.0001, **p<0.01. CBZ=carbamazepine; LEV=levetiracetam; LTG=lamotrigine; ns=not significant; WT=wild type

### SN-2000 reduces post-ictal aggression and anxiety-like behavior in wild type (WT) mice

Many first-line ASMs are associated with increased aggression or irritability, resulting in challenging clinical trade-offs between improved seizure control and psychiatric stability (Samanta 2025). While there was no difference in the time to recover from MES in mice exposed to SN-2000 60 mg/kg, SN-2000 80 mg/kg and VPA, latency to attack was doubled in SN-2000 80 mg/kg compared to VPA, suggesting a potential reduction in aggressive properties with SN-2000 (Figure 5a,b). Further, WT mice exposed to SN-2000 40mg/kg BID vs vehicle buried significantly less marbles (p<0.01, Figure 5c), and spent significantly more time in the center of an open field with novel object, suggestive of an anxiolytic effect with SN-2000 (p<0.001, Figure 5d). *Scn1a+/-* mice have shown inconsistent anxiety-like phenotypes compared to WT mice, we therefore did not include *Scn1a+/-* groups in the experimental design (Bahceci et al. 2020).

**Figure 5:**
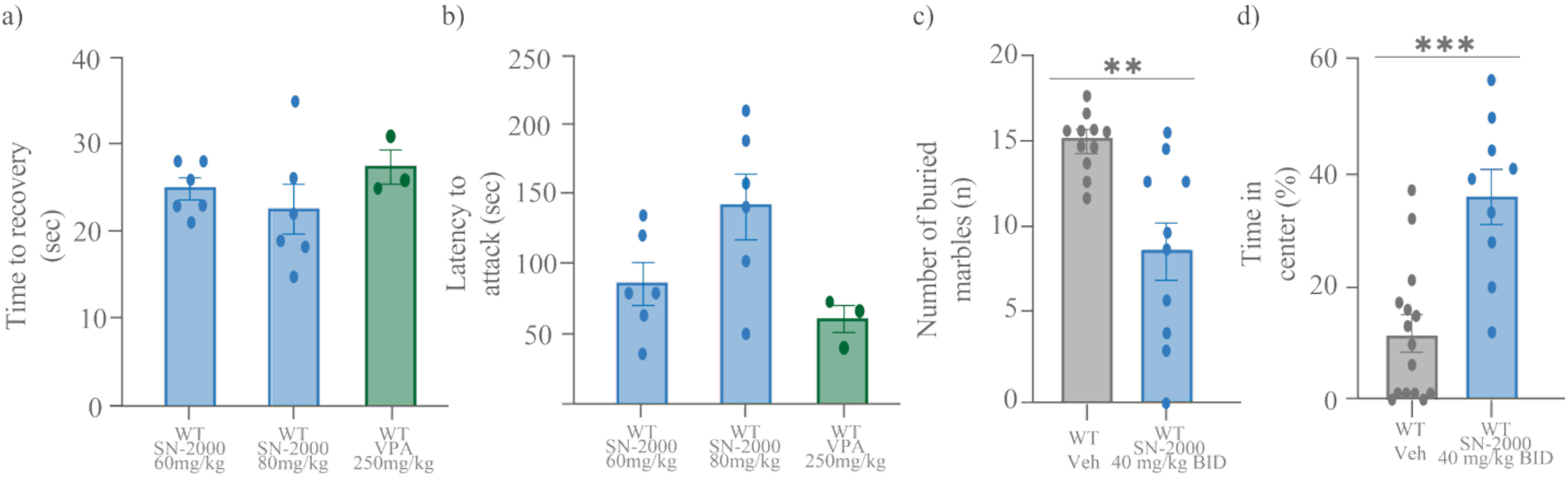
SN-2000 exposure reduces aggressive and anxiety-like behaviors in WT mice. a) Time to recovery following MES seizure induction in WT mice exposed to SN-2000 60mg/kg, SN-2000 80mg/kg, and VPA 250mg/kg. b) Latency to attack following recovery from MES seizure induction assay in WT mice exposed to SN-2000 60mg/kg, SN-2000 80mg/kg and VPA 250mg/kg. c) Number of marbles buried by WT mice exposed to SN-2000 40mg/kg BID or vehicle. d) Percentage of time in center of open field with novel object in WT mice exposed to SN-2000 vs vehicle. Unpaired t-test, **p<0.01; ***p<0.001.

### SN-2000 improves cognition in refractory epileptic and WT mice

Cognitive impairment is a common symptom of epilepsy, which may result from manifestations from the disease, from side effects of ASMs (e.g. brain fog), or a combination of both (Samanta 2025). Given the clinically substantiated link between PDE4i and improved cognition, we evaluated potential hippocampal-dependent learning and memory in *Scn1a+/-* models and WT mice. While no difference was observed in novel Y maze arm entries in WT mice exposed to SN-2000 vs vehicle, a significantly higher number of novel Y maze arm entries was observed in *Scn1a+/-* mice exposed to SN-2000 40 mg/kg BID vs vehicle (p<0.05, Figure 6a). Furthermore, increased time spent near Barnes Maze target during a probe trial was observed in both WT and *Scn1a+/-* mice exposed to SN-2000 40 mg/kg vs vehicle, indicative of potential pro-cognitive and long-term memory forming effects (Figure 6b,c). Together, these data point to a role for SN-2000 in improving cognitive function and reasoning in epileptic models.

**Figure 6:**
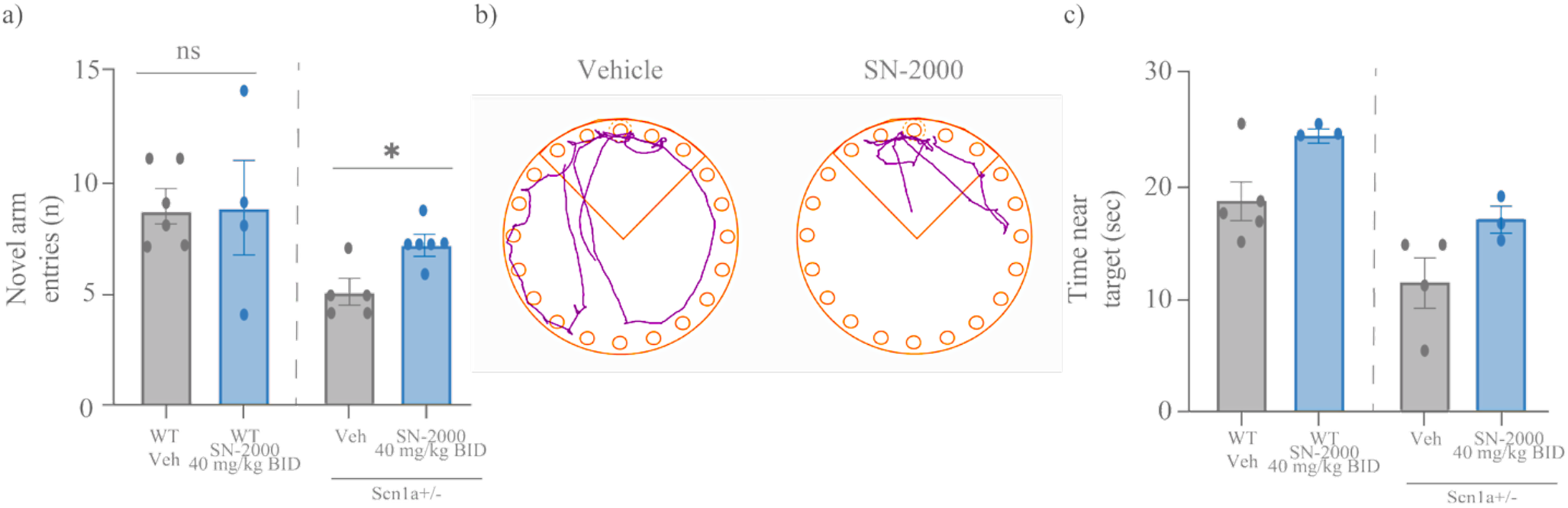
SN-2000 exposure improves memory and cognition in WT and refractory epileptic mice. a) Number of novel arm entries in forced alternating Y-maze in WT or *Scn1a+/-* mice exposed to SN-2000 40mg/kg BID or vehicle. b) Representative traces of target-seeking activity during probe trial of Barnes Maze assay in mice exposed to SN-2000 40 mg/kg or vehicle. c) Percent of time near target in Barnes Maze probe trial in WT and *Scn1a+/-* mice exposed to SN-2000 40mg/kg BID or vehicle. Unpaired t-test p<0.05

## DISCUSSION

Here we present data supporting phosphodiesterase 4 (PDE4) inhibition as a strategy for seizure reduction in a refractory epilepsy model, and describe proof-of-concept findings for SN-2000, a first-in-class, brain-penetrant, allosteric PDE4B inhibitor (PDE4Bi) with psychiatric-neutral and potential pro-cognitive properties. Treatment with SN-2000 reduced seizure activity across multiple genetic and acquired epilepsy models, with efficacy comparable to standard of care ASMs. We propose that seizure reduction is mediated by targeted elevation of cAMP in PDE4B-expressing brain regions, leading to a paradoxical regulation of excitatory and inhibitory neuronal activity, as well as reinforcement of neuronal plasticity, survival, and anti-inflammatory processes. SN-2000 also reduced anxiety-like behavior in WT mice and improved certain cognitive parameters in both WT and epileptic mice.

In a highly refractory *Scn1a+/-* mouse model, the pan-PDE4i crisaborole, a topically approved agent for atopic dermatitis (Pfizer Inc, 2016), provided partial protection from hyperthermia-induced seizures and reduced spontaneous recurrent seizures. In contrast, rolipram and roflumilast, commercially available pan-PDE4 inhibitors that were previously trialed in neurological disorders, failed to protect against hyperthermia-induced seizures at tolerated doses. This discrepancy may be attributed to the low brain penetrance of crisaborole, which allows a broader therapeutic window relative to highly brain-penetrant PDE4 inhibitors like rolipram and roflumilast, which may cause rapid supraphysiological cAMP accumulation and decreased tolerability (Burgin et al., 2010). Similarly, amlexanox, a moderately brain-penetrant pan-PDE4 inhibitor, reduced pilocarpine-induced seizures in mice as well as post-ictal elevations of hippocampal PDE4B expression (Yang et al., 2024; Liu et al., 2017). Our zebrafish data further support this mechanism, as PDE4B knockdown alone was sufficient to reduce seizure-like activity.

To achieve PDE4B isoform specificity, SN-2000 was de novo designed as the first isozyme-specific allosteric PDE4B modulator. While other PDE4B-selective inhibitors under development for peripheral disorders target the catalytic domain (NCT05321082, NCT05321069, NCT06238622, NCT05469464, NCT04982432, NCT05375955), SN-2000’s allosteric mechanism confers greater isoform selectivity, preserves physiological cAMP levels, and improves target engagement—advantages particularly important for neurological applications (Li et al., 2018). This mirrors the strategy used to develop zatomilast, an allosteric PDE4D inhibitor in clinical trials for FXS (Gurney et al., 2019).

SN-2000 was benchmarked against standard of care ASMs (VPA, CLO, CBZ, LEV, LTG, VPA+CLO), which performed in line with previous studies across *kcna1a-/-, scn1lab-/-, Scn1a+/-* (hyperthermia induction), and MES models (Baraban et al. 2013, Dogra et al., 2023; Löscher and White, 2023). SN-2000 reduced seizure activity across both teleost and murine models with comparable magnitude to these ASMs. Notably, SN-2000 doubled post-ictal latency to aggression compared to VPA, suggesting improved behavioral outcomes. VPA and other ASMs are associated with post-exposure aggressive behaviors in animal models (Bath & Pimentel, 2017; Sailer et al., 2019; Norton et al., 2020; Erath et al., 2020; Kirby-Madden et al., 2024). The anxiolytic effect of SN-2000 in WT mice is consistent with literature showing that PDE4B depletion is associated with increased anxiety-like behavior (Zhang et al., 2008). In humans, psychiatric and mood-related adverse events—such as aggression, agitation, and suicidal ideation—remain a major clinical challenge with many ASMs (Samanta et al., 2025). If clinically translatable, SN-2000 may offer seizure control with a reduced psychiatric burden.

There is a strong clinical precedent for PDE4 inhibition as a cognitive enhancer (Zeller et al., 1984; Fleischhacker et al., 1992; Hennenlotter et al., 1989; Horowski et al., 1985; Duinen et al., 2018; Blokland et al., 2019), and this continues to be evaluated in ongoing clinical trials (NCT04057898, NCT05358886, NCT05367960). In our Barnes Maze analysis, SN-2000 improved hippocampal-dependent learning and long-term memory in both WT and epileptic mice, consistent with prior findings on pan- and isoform-selective PDE4 inhibitors (Barad et al., 1998; Burgin et al., 2010; Gallant et al., 2010; Huang et al., 2007; Gurney et al., 2017). No significant effect was observed in short-term spatial memory in the Y-maze analysis in WT animals, which may reflect a ceiling effect. However, SN-2000 significantly increased novel arm entries in epileptic models, suggesting rescue of spatial memory deficits via PDE4B inhibition. Barker-Haliski et al. (2016) showed that the first-line ASMs VPA and CBZ impaired cognition in naïve rodents, while there was no impact of ASMs on fully kindled animals. In clinical populations, cognitive impairment is a common comorbidity of epilepsy and a prominent side effect of certain ASMs (Samanta et al., 2025). The current findings support further exploration of allosteric PDE4B inhibition in neurological disease with pro-cognitive potential.

Mechanistically, we build on prior literature showing that cAMP elevation via PDE4 inhibition increases inhibitory tone while suppressing excitatory drive through canonical PKA, EPAC, and HCN/CNG channel signaling (Donders et al., 2024). Inhibitory neurons benefit from cAMP-induced GABAergic vesicle recycling via PKA and EPAC activation of voltage-gated calcium channels (VGCCs) and SNARE proteins, as well as increased GABA-A1R trafficking and expression via pCREB activation (Donders et al., 2024; Nakamura et al., 2015; Fabian & Hu, 2020; Podda et al., 2007). In excitatory neurons, cAMP may transiently promote glutamate release via PKA/EPAC, followed by dampened excitability through activation of CNGCs and HCN, resulting in net membrane depolarization and increased firing threshold (Ferrendelli et al., 1980; Huang & Kandel, 1996). Our observation that interictal spike amplitude is reduced in SN-2000-treated refractory epileptic mice, without a change in overall discharge frequency, supports a modulatory capacity for PDE4i in regulating aberrant network excitability without sedative effects. The role for PDE4i in promotion of neuroplasticity, neurotransmission, neuronal survival and suppressing neuroinflammation through effector proteins including BDNF, CaMK/p38, NF-kB, among others, is well established and is extensively reviewed by Donders et al. 2025. Together, these converging mechanisms support the therapeutic rationale for PDE4Bi in restoring circuit-level homeostasis in epilepsy.

Limitations of the present study include the lack of a third genetic or acquired mammalian epilepsy model beyond *Scn1a+/-* and MES, which would enhance clinical translatability. Mechanistic claims are also preliminary; while our findings support a role for cAMP signaling and pCREB activation, additional work is needed to elucidate the impact of PDE4Bi on downstream intracellular signaling cascades. The latency to attack assay was restricted to the post-ictal period, which limits interpretation of potential long-term effects of SN-2000 on reducing aggressive behaviors. Finally, anxiety and cognitive assays lacked active comparators, limiting interpretation of SN-2000’s potential preclinical superiority to existing ASMs.

Overall, the data suggest that SN-2000 may provide a differentiated therapeutic option for epilepsy, offering broad-spectrum seizure protection with reduced psychiatric side effects and potential pro-cognitive side properties. Future research is directed toward understanding preclinical safety in higher order mammals, and fulfillment of regulatory-enabling milestones necessary for IND submission and entry into first-in-human studies.

